# Chromatin interactions correlate with local transcriptional activity in *Saccharomyces cerevisiae*

**DOI:** 10.1101/021725

**Authors:** Zackary Scholl, Jianling Zhong, Alexander J. Hartemink

**Affiliations:** Program in Computational Biology and Bioinformatics, Duke University, Durham, North Carolina, 27708, USA; Department of Computer Science, Duke University, Durham, North Carolina, 27708, USA

## Abstract

Genome organization is crucial for efficiently responding to DNA damage and regulating transcription. In this study, we relate the genome organization of *Saccharomyces cerevisiae* (budding yeast) to its transcription activity by analyzing published circularized chromosome conformation capture (4C) data in conjunction with eight separate datasets describing genome-wide transcription rate or RNA polymerase II (Pol II) occupancy. We find that large chromosome segments are more likely to interact in areas that have high transcription rate or Pol II occupancy. Additionally, we find that groups of genes with similar transcription rates or similar Pol II occupancy are more likely to have higher numbers of chromosomal interactions than groups of random genes. We hypothesize that transcription localization occurs around sets of genes with similar transcription rates, and more often around genes that are highly transcribed, in order to produce more efficient transcription. Our analysis cannot discern whether gene co-localization occurs because of similar transcription rates or whether similar transcription rates are a consequence of co-localization.

## INTRODUCTION

Genome organization is crucial for gene regulation (1-3) and DNA damage response (4,5). Recent experimental evidence suggests that genome organization has a close relationship with transcription (6-10). Evidence for RNA polymerase II (Pol II) localization has been demonstrated in HeLa cells (11-14), *Bacillus subtilis* (15), mouse B lymphocytes (16), fetal liver (17), *Schizosaccharomyces pombe* (18), and *Drosophila* (19). Sites of Pol II localization, termed “transcription factories,” are found to be rich in Pol II and nascent transcripts (20-22). The clustering of Pol II into local sites is thermodynamically favorable because of a depletion attraction (23) among large particles like Pol II (24). It is possible that nearby highly transcribed genes will be drawn toward each other because of thermodynamic factors related to the concentrated transcription complexes assembled around them. However, it is still a matter of debate whether transcription factories form when chromosomes with already-associated transcriptional complexes come together—whether by thermodynamically favorable interactions between highly transcribed chromosome segments (20) or via a self-reinforcing code of histone modification (25,26)—or alternatively whether they form when chromosomes come together to organize around concentrated polymerases, pre-localized before chromosomal association (13,16).

In this study we asked whether *Saccharomyces cerevisiae* (budding yeast) exhibits transcription-related co-localization in its genome. We analyzed the correlation between genome organization (27) and either transcription rate (28-30) or Pol II occupancy (31,32). Previous studies in budding yeast found that gene expression is modulated by position in the nucleus (33-35). Moreover, genes are often confined to specific territories in the nucleus (36), and genes with similar functions occur in adjacent positions (37). Our analysis shows that interactions between large stretches of DNA are more likely to contain highly transcribed genes, or high average Pol II occupancy. In conjunction, we find that genes with similar transcription rate or Pol II occupancy are more likely to interact with each other, indicating that transcription localization is a common phenomenon in yeast, and moreover, is most frequently associated with the most highly transcribed genes.

## MATERIALS AND METHODS

A summary of the datasets used in this study is provided in Table 1. Transcription rates were determined by Miller and colleagues through dynamic transcriptome analysis (DTA) and by Pelechano and colleagues through probing nascent mRNAs with genomic run-on (GRO). Churchman and Weissman mapped locations of active Pol II by immunoprecipitating Pol II and sequencing nascent elongating transcripts (NET-Seq). The NET-Seq dataset comes from the Gene Expression Omnibus under accession GSE25107. We performed an alignment using a recent version of the S288C yeast genome, downloaded 15 June 2011, from the Saccharomyces Genome Database (http://www.yeastgenome.org/, version 64). Bowtie 0.12.7 (38) (http://bowtie-bio.sourceforge.net) was used to perform the alignment with parameters set to report only alignments from the best strata and suppressing all alignments of reads with more than five mismatches. Active Pol II occupancy was determined by normalizing the number of reads mapped to each gene by the gene’s length. Pelechano and colleagues, and Venters and Pugh, generated Pol II occupancy profiles by performing chromatin immunoprecipitation with antibodies against different subunits of RNA polymerase II, followed by hybridization to a microarray (ChIP-chip). Three-dimensional genomic locus positions were determined by Duan and colleagues by amending the chromosome conformation capture (3C) protocol (39) with a circularization step to produce unbiased, high quality, interaction pairs. The datasets we selected are comparable because the studies all used log-phase, wild-type budding yeast cells from strain BY4741 grown in YPD (except Miller et al. grew cells in SD).

Software to produce a physical representation of a genome from chromosome conformation capture data has been recently developed by Duan et al. (http://noble.gs.washington.edu/proj/yeast-architecture/sup.html). We used the interaction map produced by this software, published in Duan et al., as well as 17 additional maps that we generated on our own using the downloaded software (because each map converges to a local minimum, we wanted to assess how robust the results were to different maps and different numbers of maps). This software determines a 3D map of the budding yeast genome by minimizing the difference between measured distance and expected distance for all pairs of locations in the genome. The expected distance is determined by assuming a standard ruler for the chromatin fibers. Although these maps are certainly not an exact physical representation of the genome, they incorporate information about polymer characteristics, interaction frequencies, and the nucleolus position in yeast, which provides the raw interaction data a useful context vis-à-vis the physical yeast genome. This “contextual” perspective supplements the raw interaction data with spatial and physical constraints that transform the raw data into a more realistic description of genome organization by determining configurations for the chromosomes that are informed by both raw interaction frequencies and spatial considerations of the chromatin fibers.

**Table 1.** Summary of datasets used in this study.

| **Data type** (*Pol II subunit*) | **Method** | **Author** |
| --- | --- | --- |
| **Transcription rate** | Dynamic transcriptome analysis (DTA) | Miller et al., 2011 |
| **Transcription rate** | Genomic run-on (GRO) | Pelechano et al., 2010 |
| **Active Pol II occupancy** (*Rpb3*) | Native elongating transcript sequencing (NET-seq) | Churchman and Weissman, 2011 |
| **Pol II occupancy** (*Rpb1,8WG16,Ser5P*) | Chromatin immunoprecipitation with hybridization to microarray (ChIP-chip) | Pelechano et al., 2009 |
| **Pol II occupancy** (*Rpo21,Rpb3*) | Chromatin immunoprecipitation with hybridization to microarray (ChIP-chip) | Venters and Pugh, 2009 |
| **Physical genomic locations** | Circularized chromosome conformation capture (4C) | Duan et al., 2010 |

Genes within the interaction map were stratified into groups according to the number of chromosomes that intersected within a sphere of a given diameter around a gene (Figure 1). The sphere’s diameter was chosen to vary across a sensible range: from being so small that it always included only one chromosome, to being so large that it never included only one chromosome. For the group of genes within a given sphere, we determined the mean transcription rate and mean Pol II occupancy. For each sphere, we also calculated a percentile based on the number of groups with the same number of genes that have a mean transcription rate, or mean Pol II occupancy, lower than the group in the given sphere. This was done by bootstrapping: sampling groups of genes from the original dataset 10,000 times, without replacement, to create a distribution of the means. Each percentile was added to a vector belonging to one-chromosome or multiple-chromosome spheres and the set of genes in that sphere were removed from further analysis to prevent multiple contributions from a single gene. A group of genes in a sphere that contained fewer than 90% of the genes after removal was not used. The vector of percentiles from the combined 18 interaction maps, across five different radii, was used to compute a kernel density profile, normalized by the number of percentiles, shown in Figure 3. We recomputed the distribution for 1, 6, 12, and 18 interaction maps and found that convergence was reached after using around 6 interaction maps (Supplementary Figure 1). Scripts to perform analysis were written in Perl and Matlab (MATLAB version 9. Natick, Massachusetts: The MathWorks Inc., 2003) and are available upon request. Figures were generated using R (R Foundation for Statistical Computing, Vienna, Austria).

**Figure 1.**
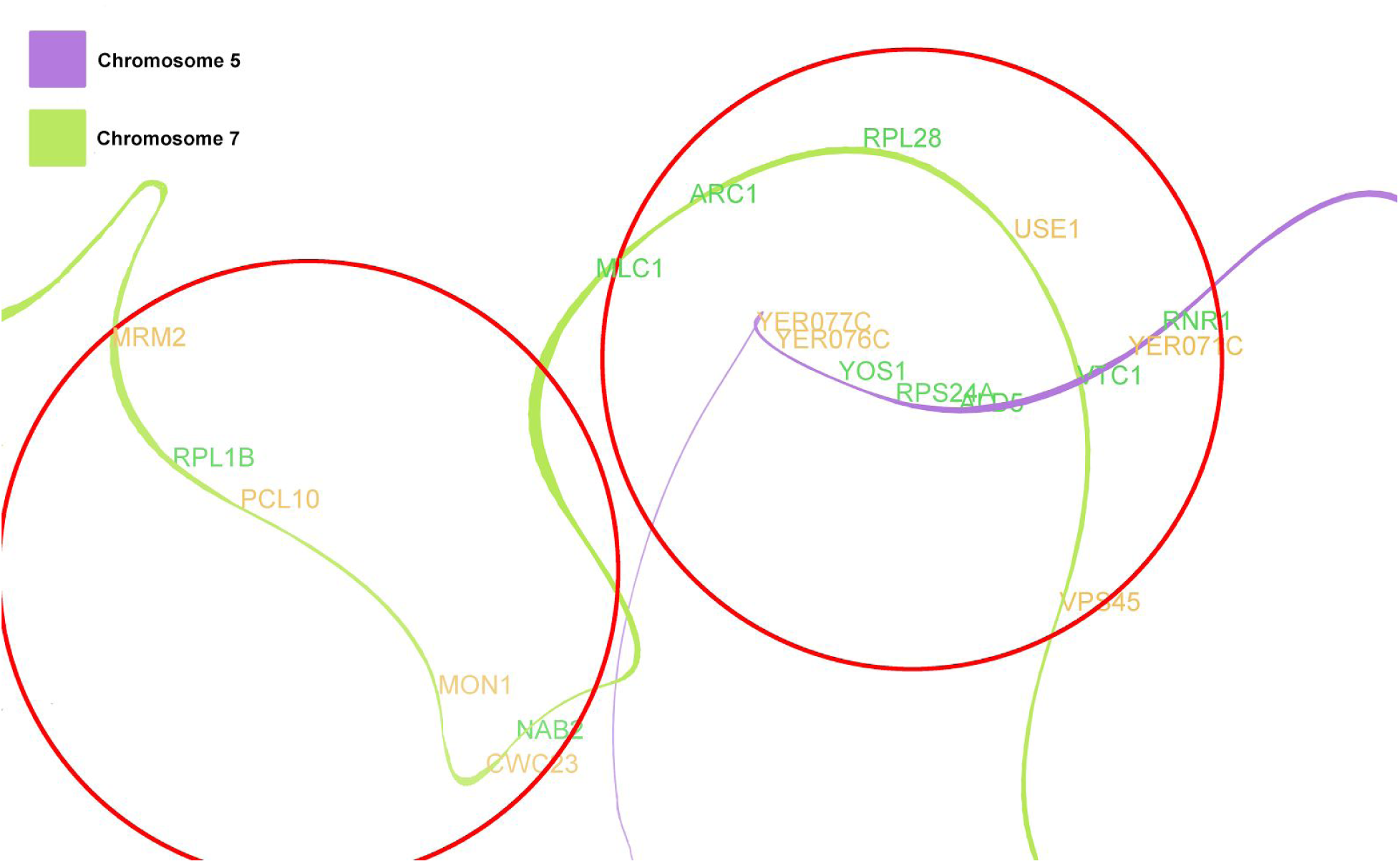
Example of contextual analysis. This shows a section of one of the interaction maps, with chromosome 5 in purple and chromosome 7 in green. Interactions are determined by stratifying the number of chromosomes that intersect a sphere of varying radius; here red circles indicate representative spheres with a diameter of 10. The gene names show the respective positions of a sample of genes, highly-transcribed genes in green and lowly-transcribed genes in orange (based on being at least one standard deviation above/below mean). The sphere on the left contains only a single chromosome and a total of 31 genes and the sphere on the right contains two chromosomes and a total of 33 genes. The percentile for a group in a sphere is determined by calculating the percent of random groups of the same size that have a lower mean than the group in the sphere. For example, from the Venters and Pugh Pol II Rpb3 dataset, the one-chromosome sphere on the left is in the 35^th^ percentile while the two-chromosome sphere on the right is in the 81^st^ percentile. This can be explained by the right sphere’s relative abundance of essential genes, like ribosome proteins, which tend to be high in Pol II occupancy.

We conducted a rank-order analysis to determine whether the genes with the highest (or lowest) transcription rate or Pol II occupancy contained more interactions (inter-chromosomal and intra-chromosomal) than random, similar to the analysis of Tanizawa and colleagues. Genes in each dataset were sorted from highest to lowest and segmented into 100 groups (these contained between 45 and 57 genes). The total number of unique gene-gene inter-chromosomal and intra-chromosomal interactions were computed for each group, based on the number of genes in that group sharing an interaction according to the 4C data (HindIII fragments at a False Discovery Rate (FDR) of 1%). This number was divided by the total number of possible interactions to get the interaction percentage between genes:

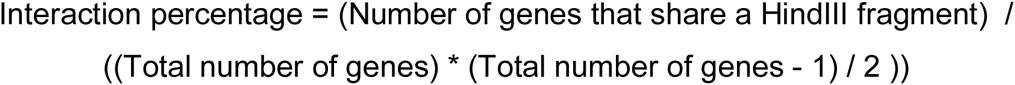

A null distribution of 90,000 interaction percentages was generated for random groups of genes of the same size (Supplementary Figure 2). The percentile (as described previously) of the interaction percentage was determined against this null distribution. One potential limitation of our method relates to the quality of the input data: the transcription rate and Pol II datasets only contain information for about 72% of the genes on average, so the rank-order analysis may be missing genes that have transcription levels or Pol II occupancy below detection level in the respective studies.

## RESULTS

### Expression in small genomic segments does not correlate with number of interactions

*Transcription rate and Pol II levels do not correlate with interaction frequency in small genomic segments.* First, we assessed the correlation between interaction frequency and expression in the absence of “contextual” information (using raw interaction frequencies but not 3D interaction maps). The correlation was computed by binning the genome into small 20kb loci, each containing about 11 genes. For each locus, we plotted the maximum transcription rate or Pol II occupancy level against the total number of inter-chromosomal and intra-chromosomal interactions. We expected that loci that interact more with other regions would have higher transcriptional activity. However, for each dataset, the comparison between interaction frequencies and transcriptional activities shows no correlation (Spearman and Pearson correlation both < 0.1; Figure 2 and Supplementary Figure 5). Since chromatin interactions exhibit some stochasticity and since the raw data are inherently noisy, this may not give an accurate description of the chromatin interactions. For example, one segment could interact often with various other loci, yet may not interact stably with any one in particular. Therefore, we suspect contextual information (polymer characteristics of chromosomes, the interactions of neighbouring loci) may be crucial for accurately depicting whether genes interact.

**Figure 2.**
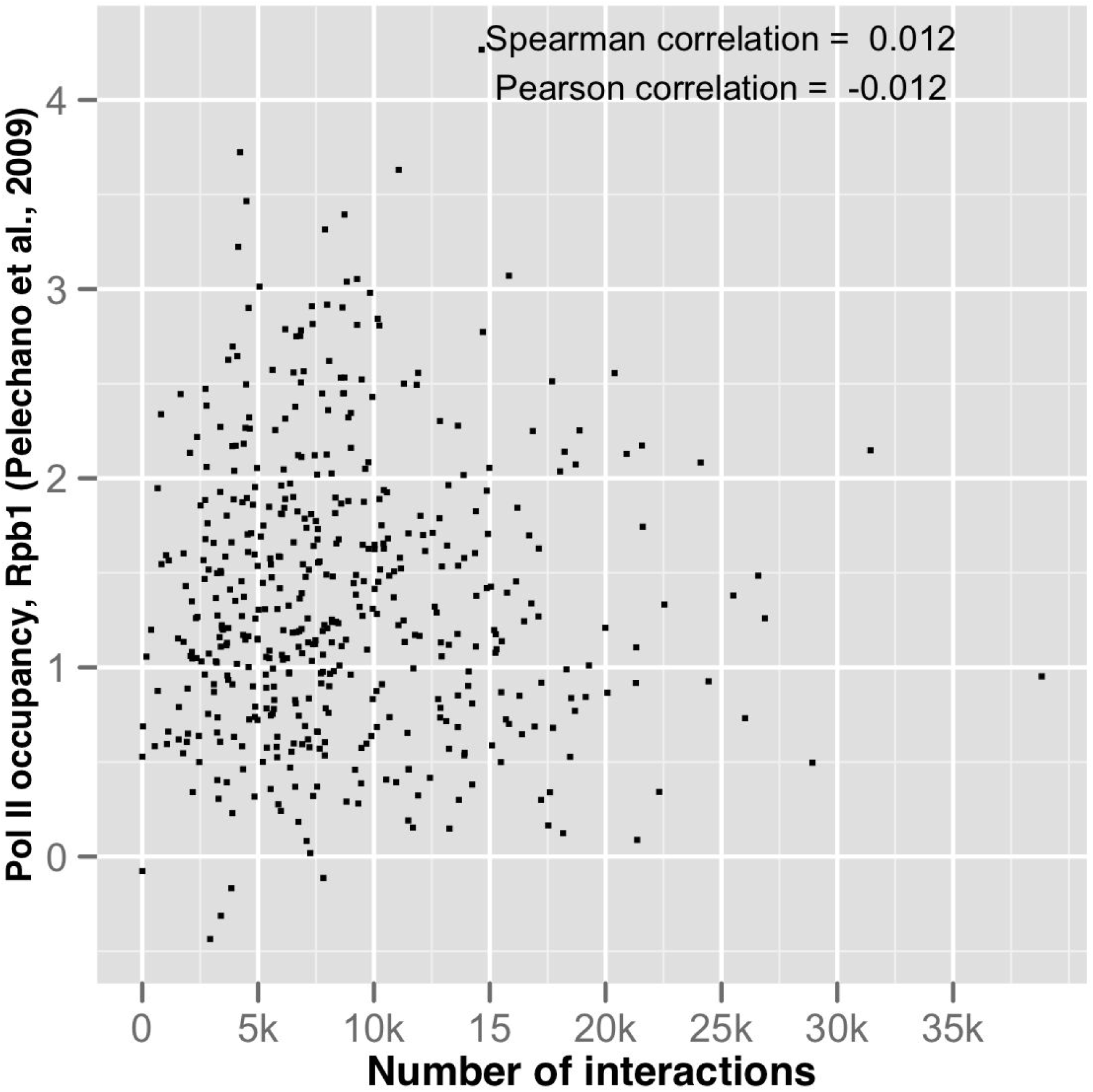
Representative example of the comparison between max Pol II occupancy of a single 25kb locus versus the number of interactions with that locus, given by the number of reads mapped to that locus from the 4C HindIII data at 1% FDR. For all the datasets compared (others are shown in Supplementary Figure 5), no correlation was found between the max transcription rate, or max Pol II occupancy, of a single locus and the total number of interactions with that locus.

### Contextual analysis reveals that interactions are correlated with expression

*Highly interacting regions of the genome are more likely to have higher transcription rates and Pol II occupancy than regions that do not interact as frequently.* This analysis uses contextual information about the genome by creating an interaction map of the genome from the raw 4C data. The interaction map is a three dimensional representation of the genome generated using interactions from the raw data. In this way, the 4C data is put into context by also including information about the polymer characteristics of chromosomes. Moreover, since all 4C interactions are accounted for simultaneously, the proximity of chromosomes in the map is likely to be dominated by stable clustering of chromosomes (meaning the chromatin regions interact frequently in the cell population), rather than one chromosome interacting with many others (meaning the interaction is transient and appears in only a small number of cells).

Using this map, we counted the number of chromosomes that were within a given distance around a small area of the genome. Areas that had no nearby chromosomes (one chromosome within the sphere) are assumed to have few interactions. Areas that had nearby chromosomes (two or more chromosomes within the sphere) are evaluated separately as they are assumed to be areas that may have more interactions, if the clustering of chromosomes is stable across the cell population. The percentile of each area was calculated for all the datasets and plotted for both single- and multiple-chromosome cases (Figure 3; other individual datasets in Supplementary Figure 3).

**Figure 3.**
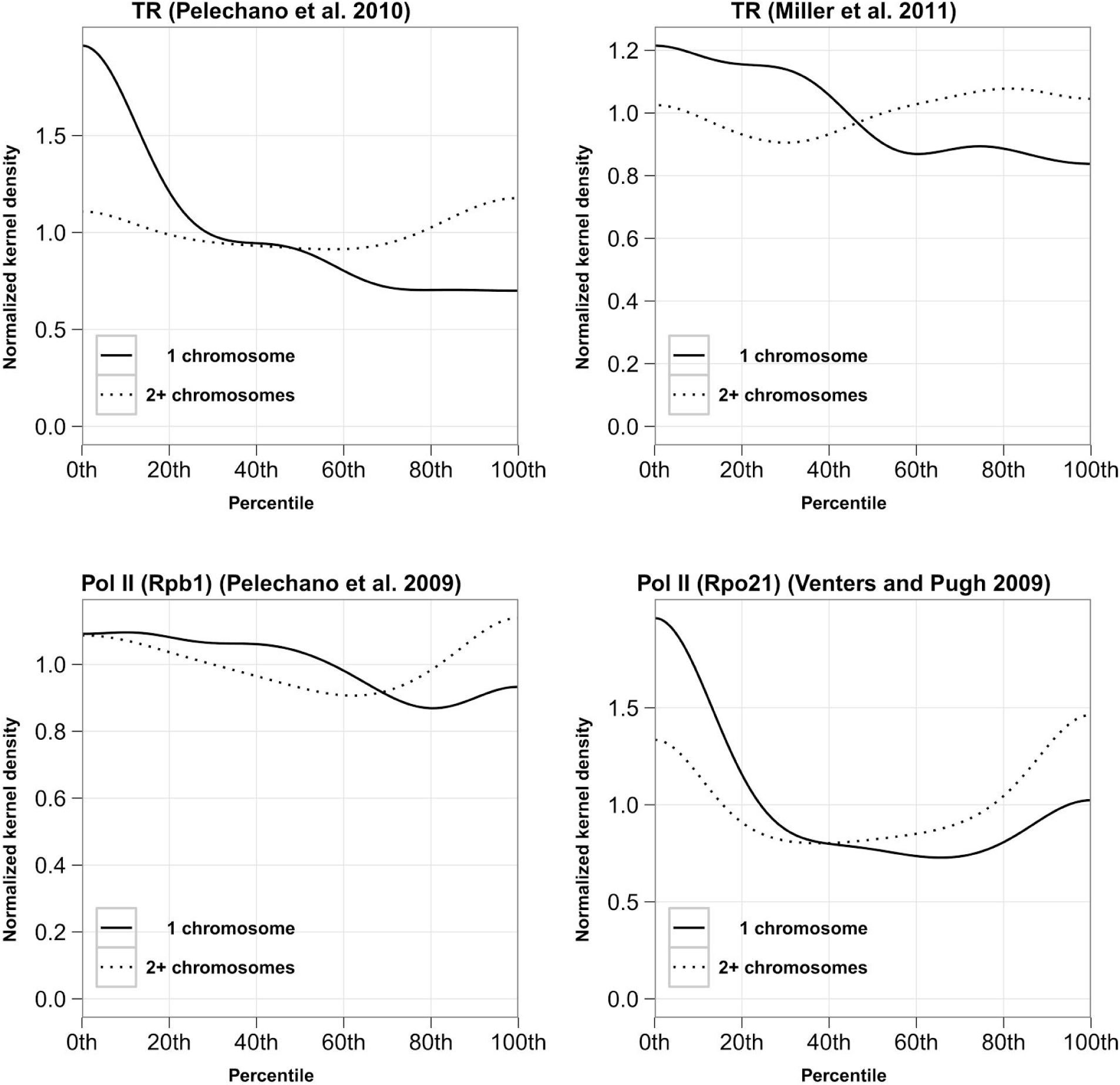
Spheres of the interaction map that have multiple interacting chromosomes (dashed lines) are more likely to have a high transcription rate than spheres with only one chromosome (solid lines). This trend emerges for both transcription rate (TR) data and RNA polymerase II (Pol II) occupancy data for various subunits of Pol II. Plots for the four other datasets are shown in Supplementary Figure 3.

For all datasets, spheres containing one chromosome (fewer interactions) are less likely to have high transcription rates and less likely to have high Pol II occupancy. For the most part, spheres containing multiple chromosomes (more interactions) have a greater likelihood of higher transcription rate and higher Pol II occupancy above the 55th percentile than spheres containing only a single chromosome. To ensure this was not an artefact of the clustering of chromosomes in a Rabl configuration near the centromere, we generated similar plots after removing genes within 50kb of the centromere. The trends of the datasets were unaffected after removing genes around the centromere (Supplementary Figure 4).

### Groups of genes with similar transcription rate or Pol II occupancy are more likely to have many interactions

*Interactions between genes with similar transcription rate or Pol II occupancy.* In addition, we examined whether groups of genes, ordered by their transcription rate or Pol II occupancy, had more interactions than would be expected by chance. Each dataset was divided into 100 groups after ranking the genes from highest to lowest transcription rate or Pol II occupancy. The calculated percentiles (see Materials and Methods) of the interaction percentage between genes in each group, in each dataset, are not uniformly distributed (combined in Figure 4, solid line), as would be expected if the groups of genes were randomly selected (Figure 4, dashed line; see Supplementary Figure 6 for individual dataset figures). Rather, the percentile of interactions from the groups are skewed toward the right. In each dataset, genes with similar levels of transcription activity or Pol II occupancy are likely to have more interactions than groups of random genes.

**Figure 4.**
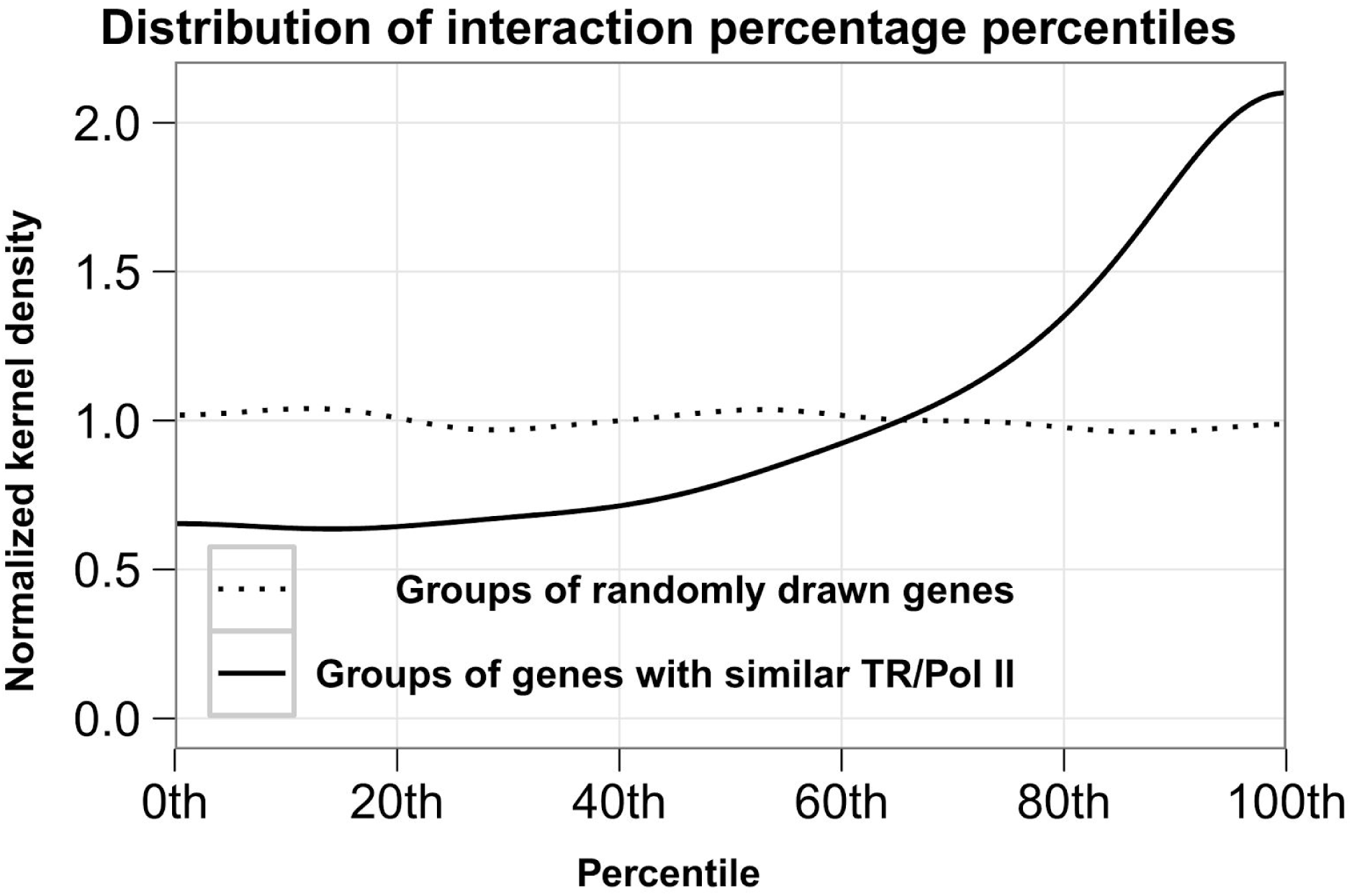
Genes were ordered according to their transcription rate or Pol II occupancy and binned into 100 groups. We computed the percentage of interactions falling into each group (number of interactions according to 4C HindIII data at 1% FDR divided by total possible number of interactions). A percentile was determined for the percentage of interaction against a null distribution of randomly chosen gene groups of the same size. This graph shows the combination of all the datasets’ distributions of interaction percentages (individual datasets are separately graphed in Supplementary Figure 6). This figure shows that genes grouped according to their transcription rate or Pol II occupancy are more likely to have more interactions (solid line) than randomly selected groups of genes (dashed line), implying that similarly transcribed or Pol II occupied genes are correlated with the number of interactions.

## DISCUSSION

In this study, we analyzed transcription and interaction data from budding yeast to determine whether interactions between genes are localized consistent with their transcription. An analysis of the raw 4C interaction data revealed no correlation between the number of interactions and the transcription rate or Pol II occupancy. However, when we used an interaction map derived from 4C data and physical information about the chromatin polymers, we find that interaction frequency is indeed correlated with transcription rate and Pol II occupancy. Though this seems contradictory at first blush, we believe evidence from the contextual analysis is more accurate because it increases the signal by incorporating more information about nuclear organization. The signal from raw interaction data in a 4C experiment is noisy because the interactions associated with a single gene could arise from many interactions with one other location, or from a few interactions with many other locations. The contextual analysis focuses on the former by concentrating on genomic locations with the most stable associations.

The accuracy of the contextual analysis is supported by looking at the percentage of interactions within groups of genes with similar transcription rates or Pol II occupancy, which uses no information about the chromosomal location. We observed that groups of genes that are similarly transcribed or Pol II occupied are more likely to have interactions within the group. Taken together, these data indicate that actual transcription localization may take place among many groups of genes of similar transcription rates to efficiently transcribe genes. The lifetime of these factories may be modulated by the relative rates of transcription needed by the group of genes, such that groups of genes with higher transcription rates may have more persistent factories. However, from this analysis it is not clear whether genes are co-localized to have similar transcription rates or whether their similar transcription rates causes co-localization. Further experiments would be needed to distinguish these scenarios.

This work provides more evidence for the transcription factory model as a general paradigm for an organism’s genomic structure. The transcription factory model suggests the clustering of Pol II, as well as the organization of chromatin around RNA polymerase clusters. One direct effect of Pol II clustering is an increase in local concentration, and thus very likely an increase in transcription initiation efficiency. This is supported by the observation that transcription initiation is generally quite slow (40), so locally concentrated Pol II could accelerate the process. However, the transcription factory model also imposes some topological and structural constraints on transcription and genome organization. Pol II complexes in clusters would be somewhat immobilized, which would imply that chromatin would have to move to be transcribed. This contradicts the canonical view of transcription, though accumulating evidence supports this new model of transcription, as reviewed in (1). Future experiments combining different cellular conditions with Pol II occupancy profiles and 3C-based data will certainly help elucidate this phenomenon.

## SUPPLEMENTARY DATA

Supplementary Data are available online: Supplementary Figures 1-6

## FUNDING

This work was supported by the National Institutes of Health [T32-GM008487-17 to Z.S.; P50-GM081883-01 to A.J.H.] and the Defense Advanced Research Projects Agency [HR0011-09-1-0040 to A.J.H.].

## ACKNOWLEDGMENTS

The authors would like to thank the organizers of the 2012 Pacific Symposium on Biocomputing for providing travel support to Z.S. and J.Z. to present portions of this work during the Structure and Function of Chromatin and Chromosomes workshop.

